# Metal speciation and bioavailability in microbial growth media

**DOI:** 10.1101/2025.10.04.680470

**Authors:** Apar Prasad, Everett L. Shock

## Abstract

Microbial growth in both natural environments and artificial media is strongly influenced by metal speciation, which can be quantitatively modeled for a given chemical composition. Despite its importance, metal speciation is rarely incorporated into the design of microbial growth experiments, often leading to misinterpretations of metal bioavailability and toxicity. In this study, we revisit two historical microbial investigations: one that drew inaccurate conclusions due to the absence of speciation calculations, and another that relied on flawed assumptions about metal speciation. Through targeted recalculations, we demonstrate how these oversights impacted the interpretation of metal–microbe interactions including the applicability of the free ion activity model (FIAM). Additionally, we perform metal speciation analyses for a bacterial growth medium to illustrate how speciation can clarify distinctions between ‘stimulatory’ and ‘non-essential’ metals. Further simulations were conducted for six DSMZ-listed microbial media and six chemical variants of a representative medium, using estimated stability constants where experimental data were unavailable. Collectively, this work underscores the value of integrating metal speciation calculations into microbial research to improve the accuracy of conclusions regarding metal bioavailability and toxicity.

## 1. Introduction

Ever since the seminal works of Sunda & Guillard 1976, it has been known that all chemical forms of a metal are not bioavailable. This knowledge is of key importance as many metals like iron and copper are essential at low concentration but toxic at higher abundance. Some of the earlier research on this topic suggested that perhaps only the free metal ion is taken up by life that gave birth to the free ion activity model (FIAM) (Morel et al. 2025). However, numerous exceptions to this model have emerged over the years (Poldoski et al. 1979, Pärt & Wikmark 1984 and Daly et al. 1990). The present scientific consensus is that in addition to the free metal ion, metal-ligand complexes are also taken up by living cells (Levina et al. 2017).

Owing to their strong interaction, metal-ligand complexes have been a topic of scientific interest for over a hundred years (Werner & Guber 1901). Many ligands like EDTA (ethylene diamine tetraacetate) and NTA (nitrilotriacetate) are regularly used as ‘chelators’ to control the bioavailability of metals in microbial growth media. Additionally, microbes have evolved to synthesize proteins called siderophores and chalkophores that selectively chelate metal ions that are essential to their biological function. The predominance of ligands in the metal distribution in these biological systems can be attributed to the high equilibrium constant for the association reaction (stability constant) between metal ions and the afore-mentioned ligands. However, this predominance is not likely to uphold for all ligands, especially at low concentration. For example, the aqueous distribution of trivalent cations like Gallium and Bismuth is heavily dominated by hydroxide complexes even in the presence of metal-chelators (Harris & Pecoraro 1983 and Sun et al. 2001). The metal distribution in a system is a complex function of all possible reactions with the metal ion and the associated stability constant and concentration.

In lieu of the complicated nature of metal-ligand interaction and metal-bioavailability, it is crucial to obtain accurate chemical distribution of metals (metal speciation) in biological systems to assess their toxicity and requirement. Direct measurement of metal speciation via analytical techniques in systems of low metal concentration with multiple ligands has not proved to be reliable (Kiss et al. 2017). Alternatively, metal speciation may be obtained theoretically under the equilibrium assumption using law of mass action and law of mass-balance. As metal-ligand complexation is quite rapid (reaching equilibrium within seconds), this is a reasonable assumption to make. These laws have been incorporated in several equilibrium speciation programs like EQ3/6 (Wolery 2010) that are routinely used to obtain elemental speciation in geochemical systems. However, these theoretical alternatives generally suffer from the lack of thermodynamic data (Kiss et al. 2017). Recently, considerable progress has been made in supplementing experimental measurements of metal-ligand thermodynamic properties with estimates of stability constant and entropy (Prasad & Shock 2025a and Prasad & Shock 2025b). This database and the estimation techniques therein permit metal speciation calculations for a large set of metal-ligand complexes from 0°C to 125°C.

In this study, we performed metal speciation calculations across a variety of growth media commonly employed in bacterial and algal cultivation, and examined their alignment with published data on metal bioavailability. To provide conceptual clarity, the following section defines the terms *metal speciation* and *metal bioavailability*, supplemented with illustrative examples and a brief discussion of their interrelationship. We then applied speciation modeling to an algal metal-toxicity study that lacked direct speciation measurements, offering new insights into the chemical environment experienced by the test organism. A similar analysis was conducted for a copper bioavailability study involving a marine amphipod, where we identified and addressed flawed assumptions regarding copper speciation. In the subsequent section, we evaluated a bacterial growth medium used in food production that reported differential requirements for manganese and zinc, and explored how speciation dynamics may account for these observations. Finally, we present speciation calculations for all defined and mineral-based media listed on the DSMZ website, highlighting how minor compositional changes can substantially influence metal speciation profiles. All stability constants used in the speciation calculations are provided in the Supplementary Table.

## 2. Metal speciation and metal bioavailability: definitions and relationship

Metal speciation is defined as the chemical distribution of the total metal in different chemical forms. For example, if 1 millimole salt ZnCl_2_(cr) is dissolved in 1 liter of pure water at 25°C and 1 bar, its speciation is given as 99.65% Zn^+2^(aq) while the remaining 0.35 % of the total zinc is as Zn(OH)^+^(aq), Zn(OH)_2_, ZnCl^+^ and ZnCl_2_(aq). Comparatively, in a 1mM aqueous solution of Zn(Oxine)_2_, 98.92% of the total zinc is as Zn(Oxine)_2_(aq), 1.06% as Zn(Oxine)^+^ and the remaining as the afore-mentioned species. Thus, metal speciation is heavily dependent on the ligands present in the system.

Metal bioavailability remains a concept with fluid and context-dependent definitions. It is commonly employed to describe those metal species capable of translocating across the lipid bilayer from the bulk solution via any cellular transport mechanism. However, its usage can be inconsistent across studies. For instance, while metal precipitates are typically excluded from bioavailable fractions, certain small, insoluble metal complexes are considered bioavailable (Schwarzenbach et al., 2003; Levina et al., 2017). Furthermore, the classification of metal–ligand complexes varies: chelates such as EDTA are generally regarded as non-bioavailable (Campbell, 1995), whereas complexes with ligands like oxine have been shown to facilitate bioavailability (Ahsanullah & Florence, 1984).

The relationship between metal speciation and bioavailability has now been observed for a diverse set of organisms like prokaryotic microorganisms (Sunda & Gillespie 1979, Sunda & Ferguson 1983 and Morton et al. 2000), eukaryotic microorganisms (Sunda & Guillard 1976, Anderson & Morel 1978, Sunda & Huntsman 1986, Sunda & Huntsman 1992 and Canterford & Canterford 1980), animals (Sunda et al. 1978 and Daly et al. 1990) and human cell cultures (Zhu et al. 2006, Scheers et al. 2014, Peng et al. 2014, Pereira et al. 2014, Nday et al. 2012 and Hart et al. 2015). While all these studies report a qualitative difference in biological response to changes in metal speciation, very few have attempted to establish quantitative relationships between metal speciation and bioavailability. Most of these studies (like Sunda & Guillard 1976 and Engel et al. 1981) report a trend like the schematic demonstrated in Fig. 1 where the biological response (like metal-toxicity) has no correlation with total metal concentration but mimics a smooth mathematical function when plotted against the concentration of the perceived bioavailable species. Such a mathematical relationship is quite desirable as not only is it founded in theory (Morel et al. 1979), it may also inform which metal species may be bioavailable and which species are unlikely bioavailable candidates. An established quantitative model of metal speciation and bioavailability can simulate experiments and predict the conditions where the presence or absence of metals has significant biological impact. While it is understood that this relationship is likely to get more complicated in more complex systems comprising of multiple bioavailable species and multiple transport pathways, efforts are nevertheless warranted owing to the enormous power that such correlations might bring. We have tried to create these correlations in the following sections after replicating one of the well-received studies on metal speciation and bioavailability that validates our stability constant data and simulation methodology (Supplementary Material S1).

**Fig. 1.**
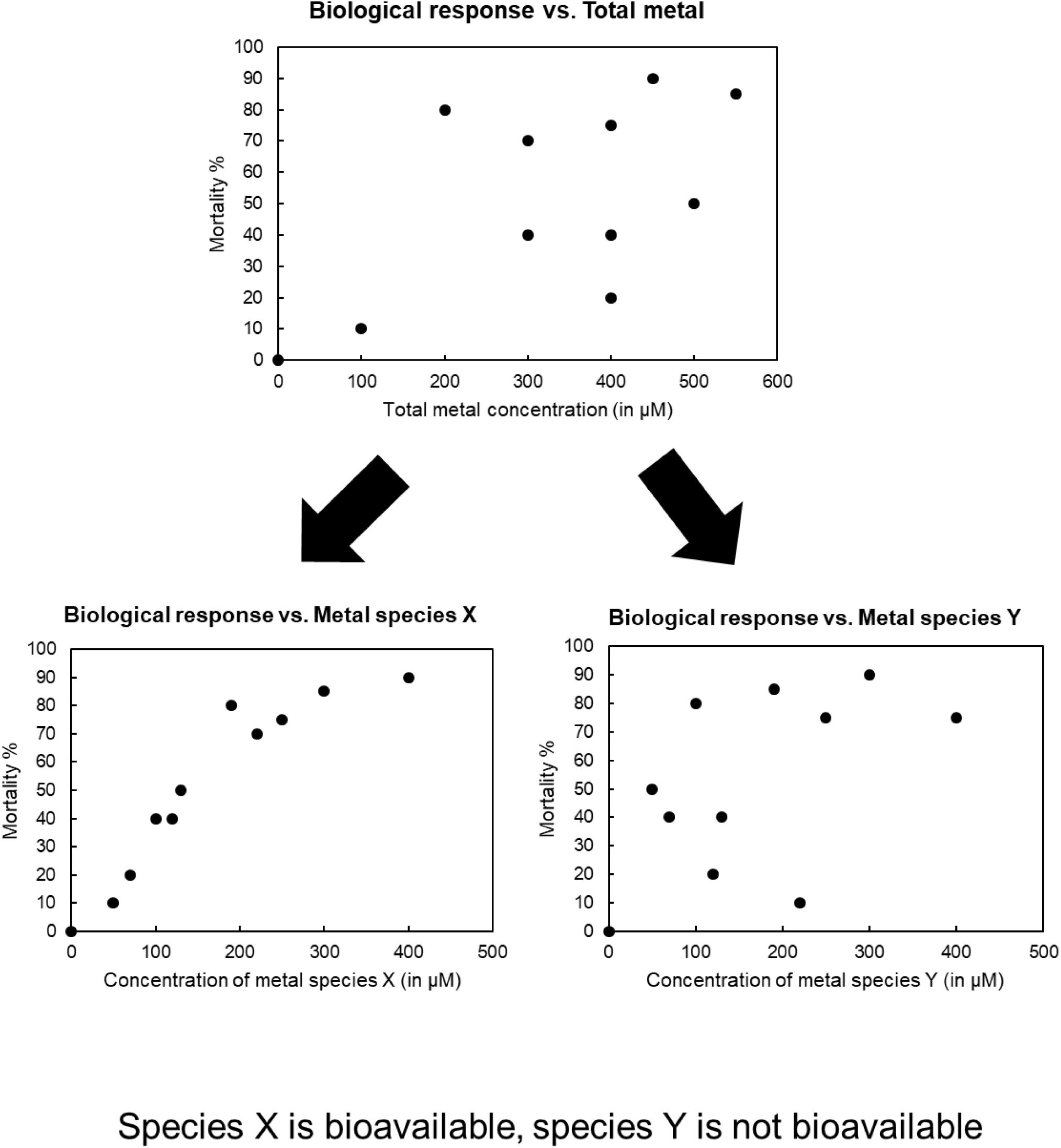
**Schematic of relationship between speciation and bioavailability.**

## 3. Simulation of a bioavailability study without speciation analysis

In 1980, Laube et al. published a study of metal toxicity on the blue-green alga (Cyanobacterium) *Anabaena*, strain 7120. In the study, Laube et al. examined the differences in the growth of the alga measured using optical density measurements over a wide range of cadmium, copper and lead concentrations (10^-8^ to 10^-3^ M) with and without equimolar concentrations of NTA in a defined growth medium over 15-20 days. Quite remarkably, Laube et al. 1980 reported that NTA ‘did not reduce, and in some cases increased, metal toxicity to *Anabaena* 7120’ and suggested that ‘these metals do not act on this alga only in the ionic form’, thereby against the free ion activity model. However, no speciation calculations were reported, nor any attempts made subsequently by studies citing the paper. One study deemed it ‘problematic’ and suggested that increased toxicity with high NTA concentrations might still be caused by appreciable concentrations of the free ions due to competition with metals like Fe or due to volumetric errors (Campbell, 1995).

To obtain a clearer understanding of this study, we performed speciation calculations for the growth media used by Laube et al. 1980 using critically evaluated thermodynamic data from the literature and estimates of metal-ligand stability constants for the unmeasured complexes (Prasad & Shock 2025a and Prasad & Shock 2025b). The calculations were performed using the software EQ3/6 (Wolery 2010) at 25°C and 1 bar as that was similar to the experimental conditions reported in the paper (20°C and supposedly 1 atm) and happen to be the conditions where most experimental thermodynamic data is reported. To investigate whether these experiments were indeed in disagreement with the free ion activity model, we obtained the activity (molality*activity coefficient) of the free metal ion for both set of experiments-with and without chelator. The calculated logarithm of the free cadmium activity (log a_Cd+2_) is given next to the respective experiments in Fig. 2a. As may be seen from the figure, log a_Cd+2_ is almost identical for both set of experiments (with and without NTA) for the sub-micromolar experiments and the millimolar experiments (i.e 10^-3^ M, 10^-6^ M, 10^-7^ M and 10^-8^M total cadmium). The similarity of the growth curves for these set of experiments along with our calculated activity of the free metal suggests that the algal growth agreed with the free ion activity model, in stark contrast to the conclusions reached by the authors of the original study. The higher growth for the experiments done without NTA at 10^-5^M Cd_T_ in comparison to 10^-6^M Cd_T_ belies explanation and perhaps is due to volumetric errors as suggested by Campbell 1995. The negligible growth for experiments performed at 10^-4^ Cd_T_ may be explained by the excessively high levels of free cadmium for both set of experiments (corresponding to high micromolar values) and thus is still not in disagreement with the free ion activity model. This example clearly illustrates how erroneous conclusions about metal bioavailability may be reached without the knowledge of metal speciation.

**Fig. 2 (a-c).**
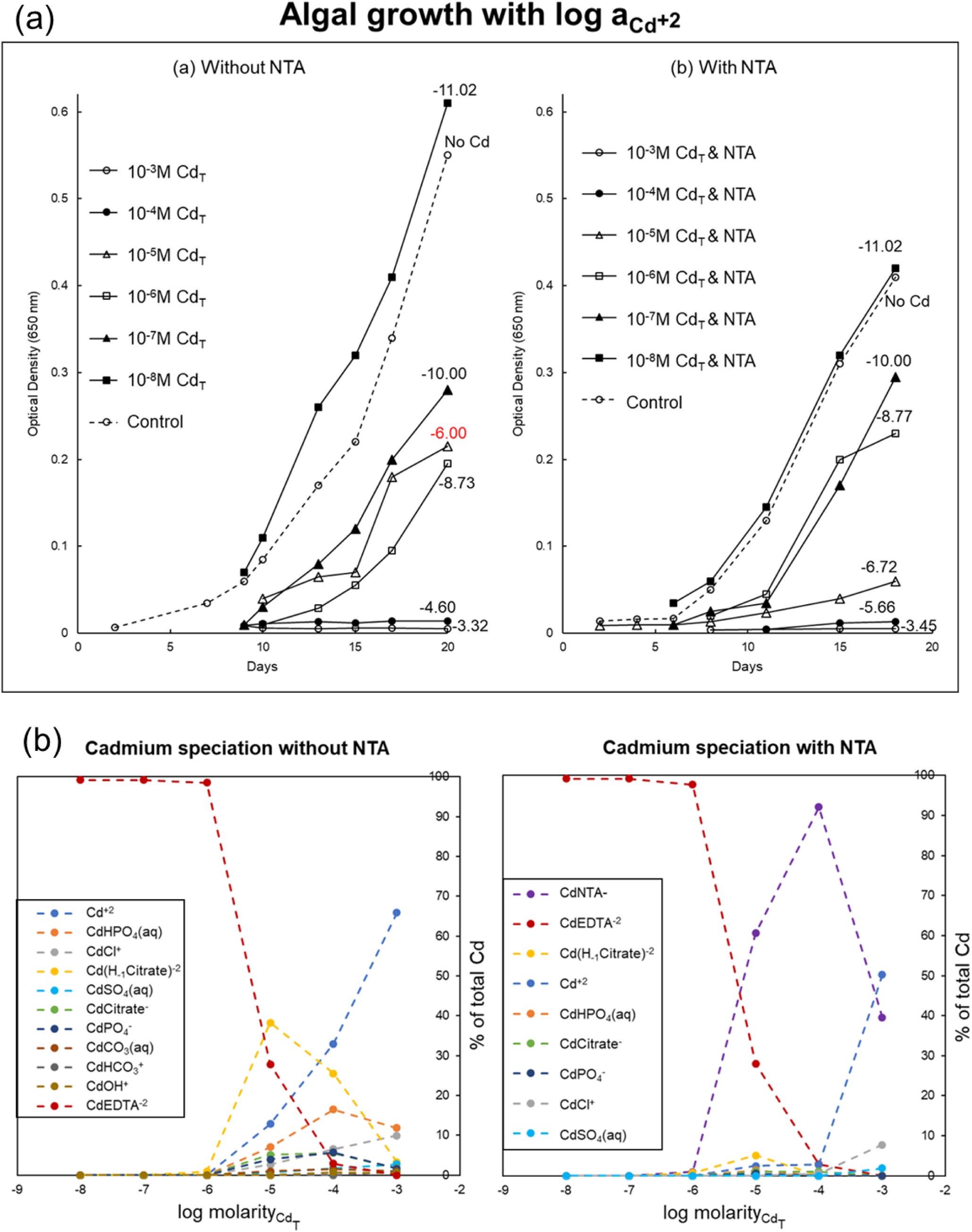
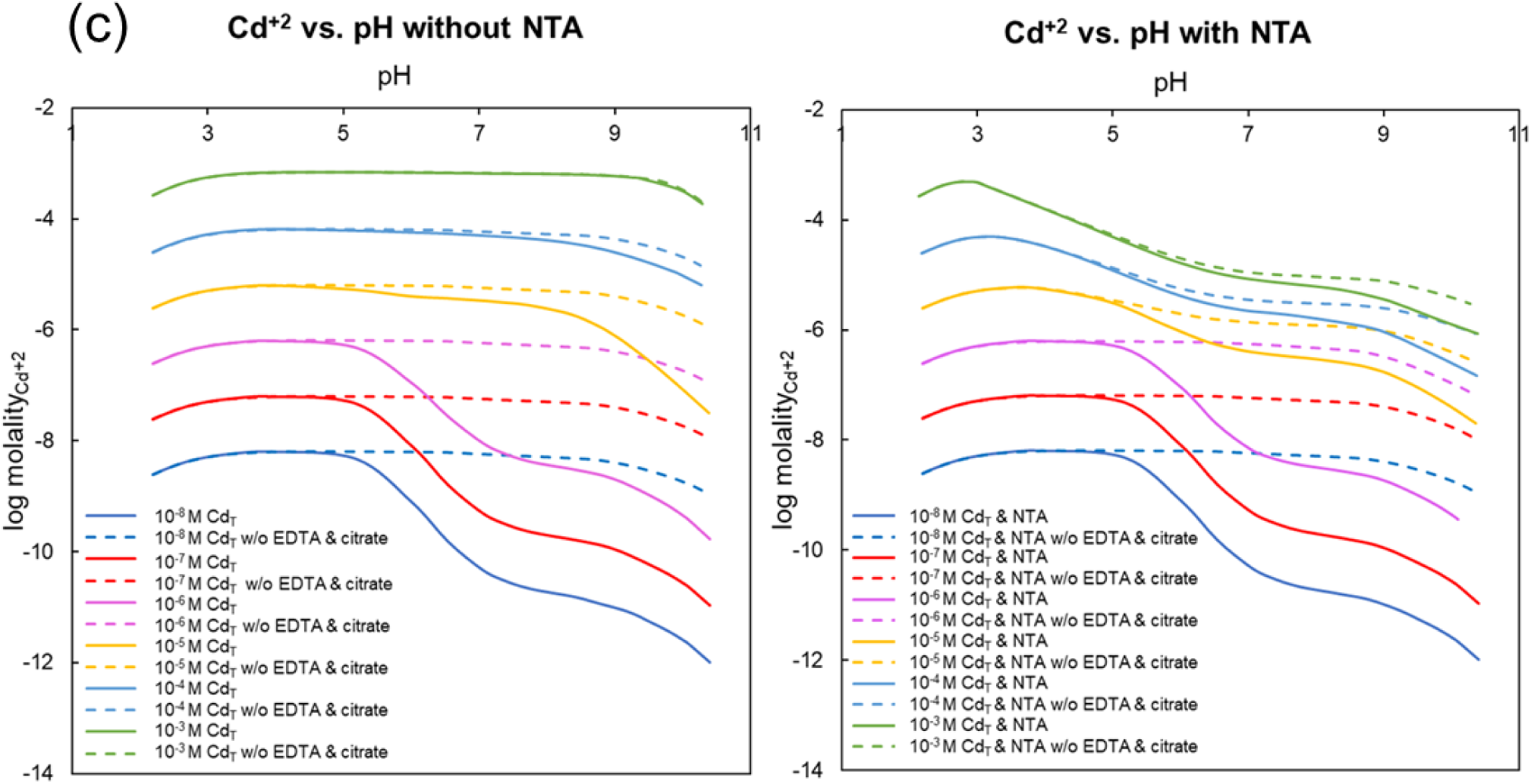
**Cadmium speciation in Laube et al. 1980.**

Perhaps the reason for the conclusions reached by Laube et al. stems from the common assumption that a chelator always dominates the metal speciation in all conditions for all metals. Our speciation calculations for cadmium (Fig. 2b) demonstrate that this is clearly not the case in these experiments. Our calculations show that for experiments with NTA, the Cd-NTA complex only dominates the speciation for the experiments carried out at 10^-4^ Cd_T_ and 10^-5^ Cd_T_. This is because cadmium speciation at sub-micromolar cadmium concentration (10^-6^M, 10^-7^M and 10^-8^M Cd_T_) is dominated by EDTA that is present in the default growth medium. As EDTA has octahedral coordination and is known to completely envelope the metal ion, the metal ion cannot interact with membrane proteins and cross the cell membrane.

While the addition of NTA in the manner of original administration does not seem to affect the free cadmium concentration, we found that this is liable to change at higher pH (Fig. 2c). For the super-micromolar concentrations of cadmium (10^-5^M, 10^-4^M and 10^-3^M Cd_T_), the free cadmium concentration by 1-3 orders of magnitude from pH 7-9. We predict that under these conditions, the addition of NTA is likely to cause a change in cadmium toxicity under the default assumption of free ion activity model. Additionally, we have presented the free cadmium concentration if these experiments are carried out in the absence of the other chelators-EDTA and citrate. As can be seen from the figure, these curves closely mimic the original curves with the chelators without being identical. In this way, speciation calculations can be used to predict bioavailability and choose the right set of conditions to verify such predictions with experiments.

Similar explanations can be given for the lead toxicity experiments made in the study. The growth curves were largely similar for experiments performed with or without NTA. However, these experiments were marred by precipitation and no account was reported of the lead lost in these precipitates. Thus, speciation calculations could not be made for these experiments to verify the compliance with the free ion activity model.

While addition of NTA did not largely affect growth for cadmium and lead experiments, it certainly decreased algal growth in the case of copper. Expectedly, the free copper activity (a_Cu_^+2^) could not explain the growth curves. Nevertheless, we also found that these growth curves could not be explained with the combination of (a_Cu_^+2^) and (a_Cu-NTA_) as suggested by Laube et al (data not shown). Thus, we could not prove that Cu-NTA complexes were bioavailable as no qualitative or quantitative relationship could be established between Cu-NTA abundance and algal growth. We believe that this may be due to a different uptake mechanism of copper as also suggested in the original study. Thus, these experiments warrant further investigation to lend a better understanding of metal bioavailability.

## 4. Simulation of a bioavailability study with erroneous speciation calculations

As the cell membrane is largely hydrophobic in nature, molecules with hydrophobic moieties like benzyl groups have a high tendency to cross the lipid bilayer via passive diffusion. This ability to cross the lipid bilayer could perhaps also be extended to metal complexes of ligands like oxine that have two electron-bearing groups (phenol and pyridine) attached to a benzene ring. To investigate this, Ahsanullah & Florence (Ahsanullah & Florence 1984) conducted a toxicity study in artificial seawater on a marine amphipod *Allorchestes compressa*, in which they analyzed copper-induced mortality with and without oxine.

Copper toxicity without oxine was carried out over a wide range of total copper concentration (black circles in Fig. 3a) in organic-free seawater (organics destroyed by ultraviolet radiation) over 96 hours. Copper toxicity experiments with oxine were carried out in the same medium with 0.5 μM, 1 μM and 2 μM oxine (black squares in Fig. 3a). Comparing the two set of experiments, Ahsanullah and Florence concluded that copper-oxine complexes (the assumed bioavailable metal species in experiments with oxine) were more toxic than free copper ion (the assumed bioavailable metal specimen in experiments without oxine) as lesser copper elicited the same or higher mortality with the addition of oxine.

**Fig. 3(a-c).**
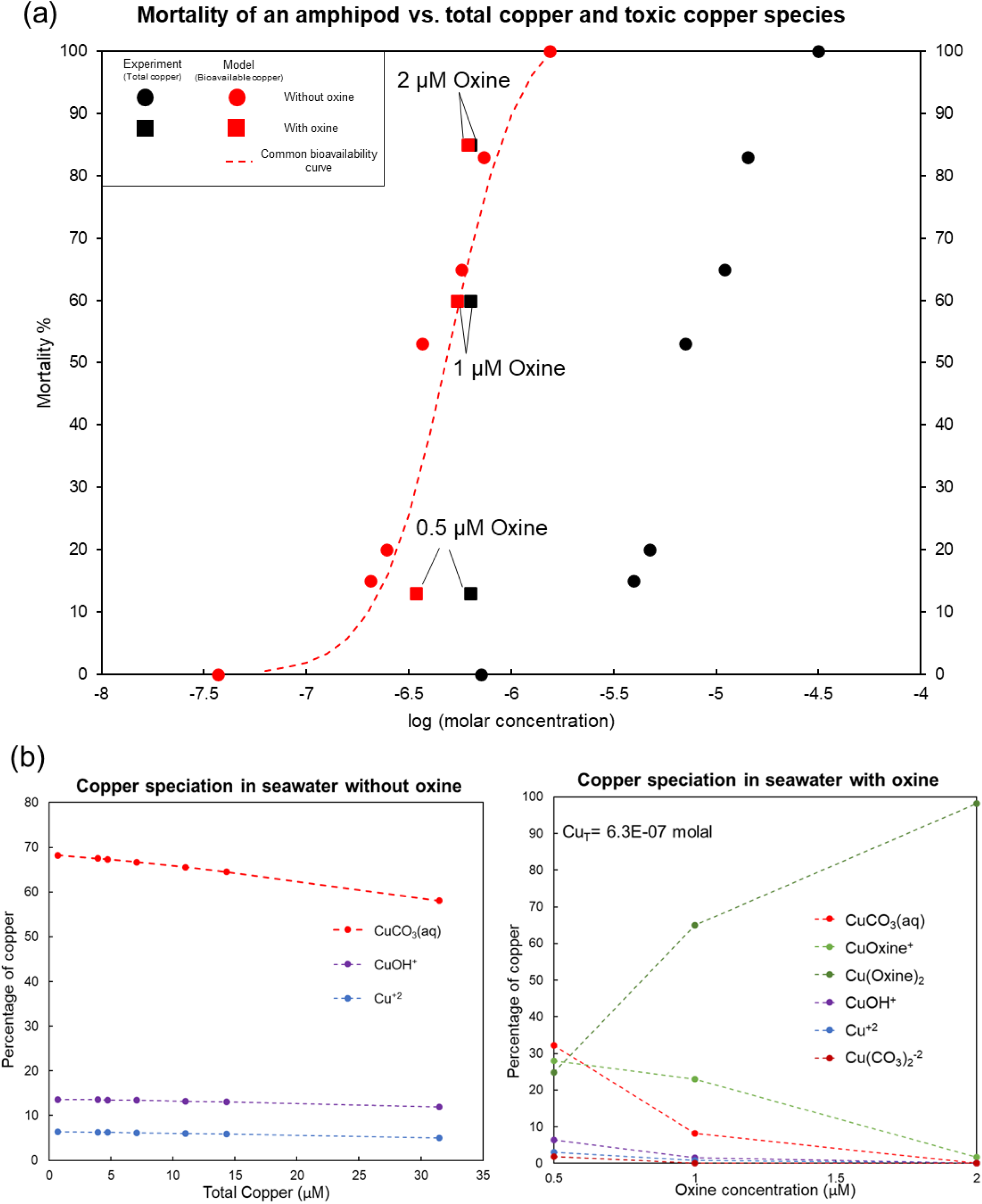
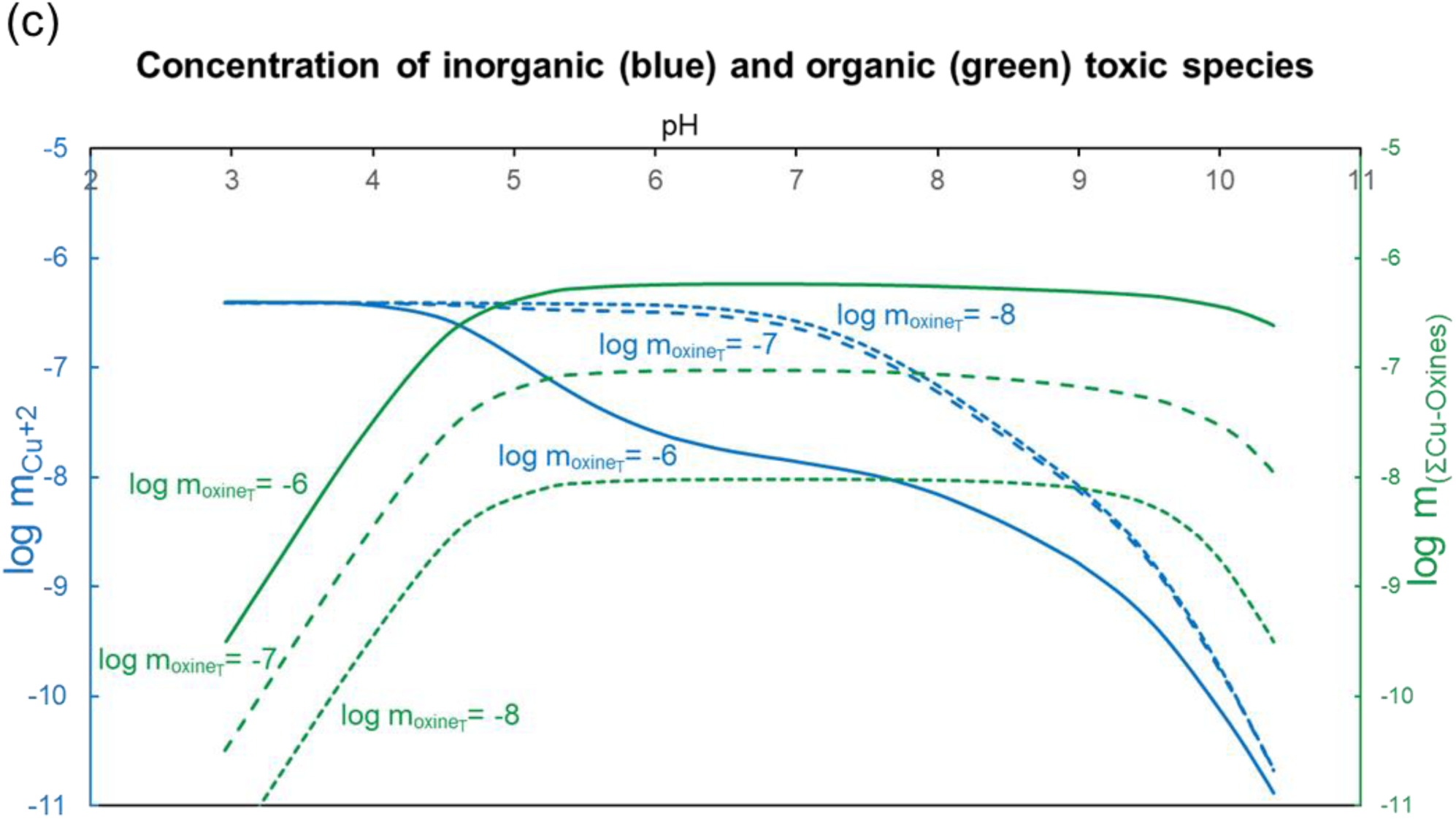
**Speciaton calculations of Ahsanullah & Florence 1984.**

However, the authors of the original study did not directly compare the mortality curves with the free copper and copper-oxine abundance as they had not performed speciation measurements or calculations for experiments performed without oxine. While they correctly obtained copper speciation for experiments with 2 μM oxine, they wrongly assumed that this speciation would hold for experiments performed at lower oxine concentration (Fig. 2b). We performed the speciation calculations for all experiments using critically evaluated metal-ligand stability constant data from the literature or using analogous estimates obtained via linear free energy relationships (Prasad & Shock 2025a and Prasad & Shock 2025b). We subsequently compared the copper-induced mortality to the abundance of actual bioavailable species-free copper ion for experiments without oxine (red circles) vs. both free copper ion and copper-oxine complexes for experiments with oxine (red squares). This assessment revealed that the biological response for free copper ion is very similar to that of copper-oxine species, in clear disagreement with the conclusion reached by the authors of the original study. Even more glaringly, free copper ion elicits a higher mortality at lower concentrations in comparison to the 0.5 μM oxine experiment.

The reason for the difference between the free copper (Cu^+2^) concentration and the total metal concentration can be seen in Fig. 3b. Only about 5-7% of the total copper is as Cu^+2^, which is perhaps why Cu^+2^ (red circles) and Cu_T_ (black circles) are separated by about 1.5 log units in Fig. 3a. Also, while our speciation calculations of copper with 2 μM oxine (Fig. 3b) matched those of Ahsanullah & Florence 1984, we found that the speciation was different at other oxine concentrations, particularly in 0.5 μM oxine where CuCO_3_ (aq) is the predominant copper species.

To predict copper toxicity in diverse environments, we performed speciation calculations in different conditions and found that copper speciation was extremely sensitive to pH (Fig. 3c). While copper-oxine complexes (green) dominate at high pH, free copper dominates the copper speciation at pH lower than 4.8 even at micromolar concentration of oxine. This may be due to the complete protonation of oxine as the pK_a_ values associated with the two ionizable groups of oxine are 9.82 and 4.92. This suggests that free copper ion dominates copper speciation at low pH and may serve as the predominant bioavailable form under these conditions.

## 5. Speciation in a bacterial growth media with contrasting transition metal requirements

*Leuconostoc mesenteroides* is a fermentative bacterium employed in several foods like kimchi and sauerkraut. In 2012, a study was published presenting an optimal growth medium for the strain ATCC8293 using the single omission technique (Kim et al. 2012). In this technique, growth is monitored as single nutrients are systematically removed from the medium. If the growth rate with the removal of a nutrient is less than 50% of the maximum growth rate, the nutrient is classified as ‘essential’. The nutrient was termed ‘stimulatory’ if the omission led to a growth rate between 50-90% and ‘non-essential’ if the growth rate was greater than 90%. Along with amino acids and vitamins, the impact of transition metals on growth was also assessed and manganese was found to stimulate growth unlike iron (II), zinc, cobalt and copper.

This led us to speculate if metal speciation could be playing a role in the differential bioavailability of transition metals. We obtained equilibrium constants from the literature or made estimates (Prasad & Shock 2025a and Prasad & Shock 2025b) for all protonation and metal-complexation reactions that could take place in the medium. Equilibrium speciation calculations were made using EQ3/6 for the composition and conditions given in the study. As manganese was a ‘stimulatory’ nutrient, it was included in the composition as a sulfate salt while the other transition metals were not part of the chemically defined minimal medium. We performed speciation calculations for the composition reported in the study and exchanged manganese with the other transition metals to simulate the respective omission experiments.

The speciation for all ligands was governed by their respective pK_a_ values as their concentration was much higher than the transition metal concentration. The speciation of the divalent transition metals in the respective omission experiment simulations was found to be case dependent. While manganese speciation was dominated by citrate, zinc and copper speciation was dominated by arginine while cobalt and iron (II) speciation was dominated by phosphate and hydroxide respectively (Fig. 4a). More specifically, manganese speciation was dominated by Mn(Citrate)^-^, zinc speciation by Zn(Arginine)^+2^, copper speciation by Cu(Arginine)^+2^, cobalt speciation by Co_2_(OH)_3_^+^ while iron (II) speciation was dominated by aqueous FeHPO_4_^0^. As small molecule transport across the cell membrane is heavily dependent on charge, it is perhaps worth deliberation if the unique requirement of manganese in the medium is because it is predominantly distributed as a negatively charged species. We additionally found that decreasing the arginine concentration by a factor of 10 substantially increased the predominance of citrate complexes for most metals. As a substantial fraction of all these metals (except iron) is present as the negatively charged citrate complex, perhaps they would have similar bioavailability. Further experimental investigation on this topic can shed more light on the speciation-bioavailability relationship and on transport of small molecules across biological membranes.

**Fig. 4(a-c).**
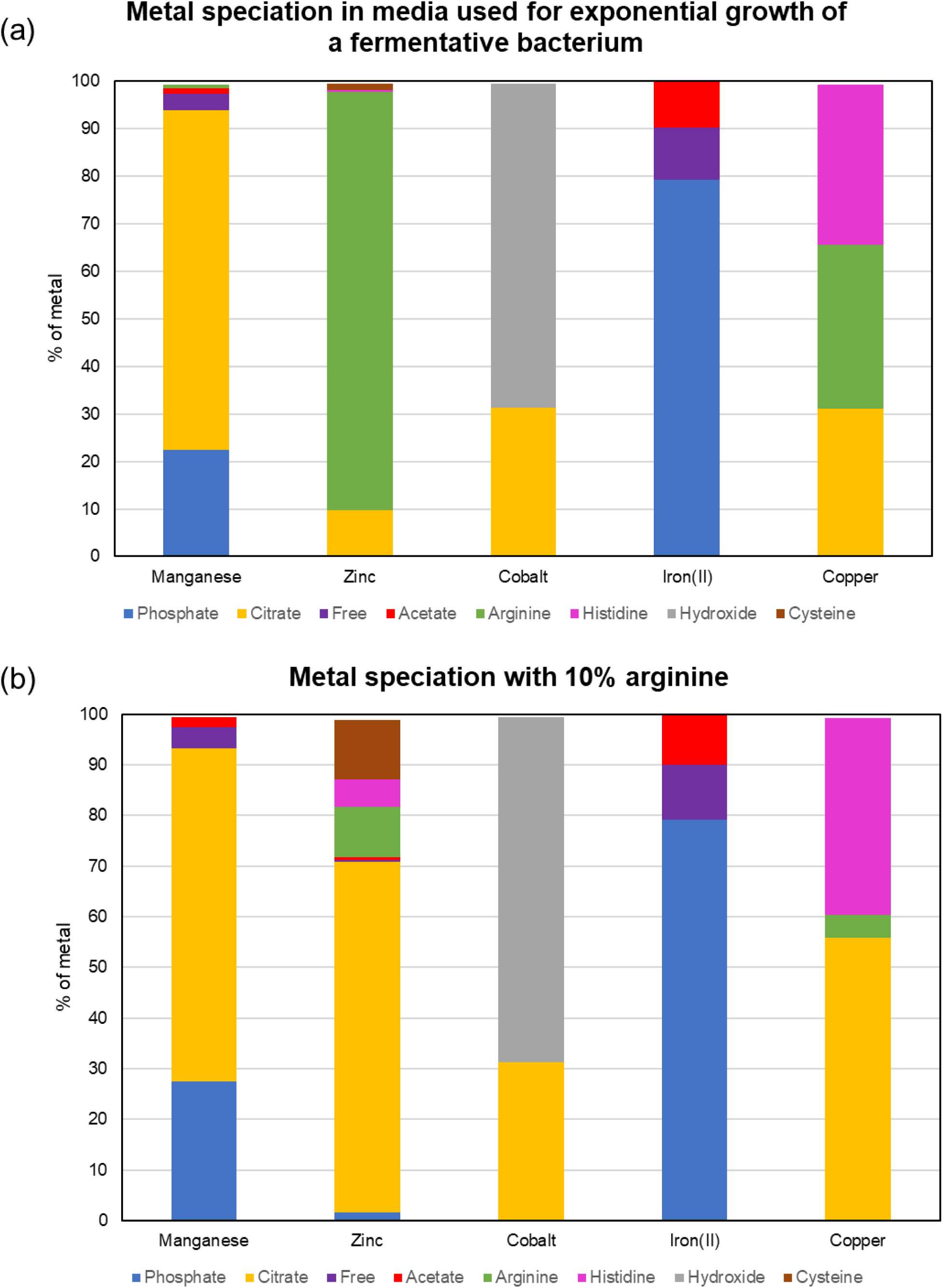
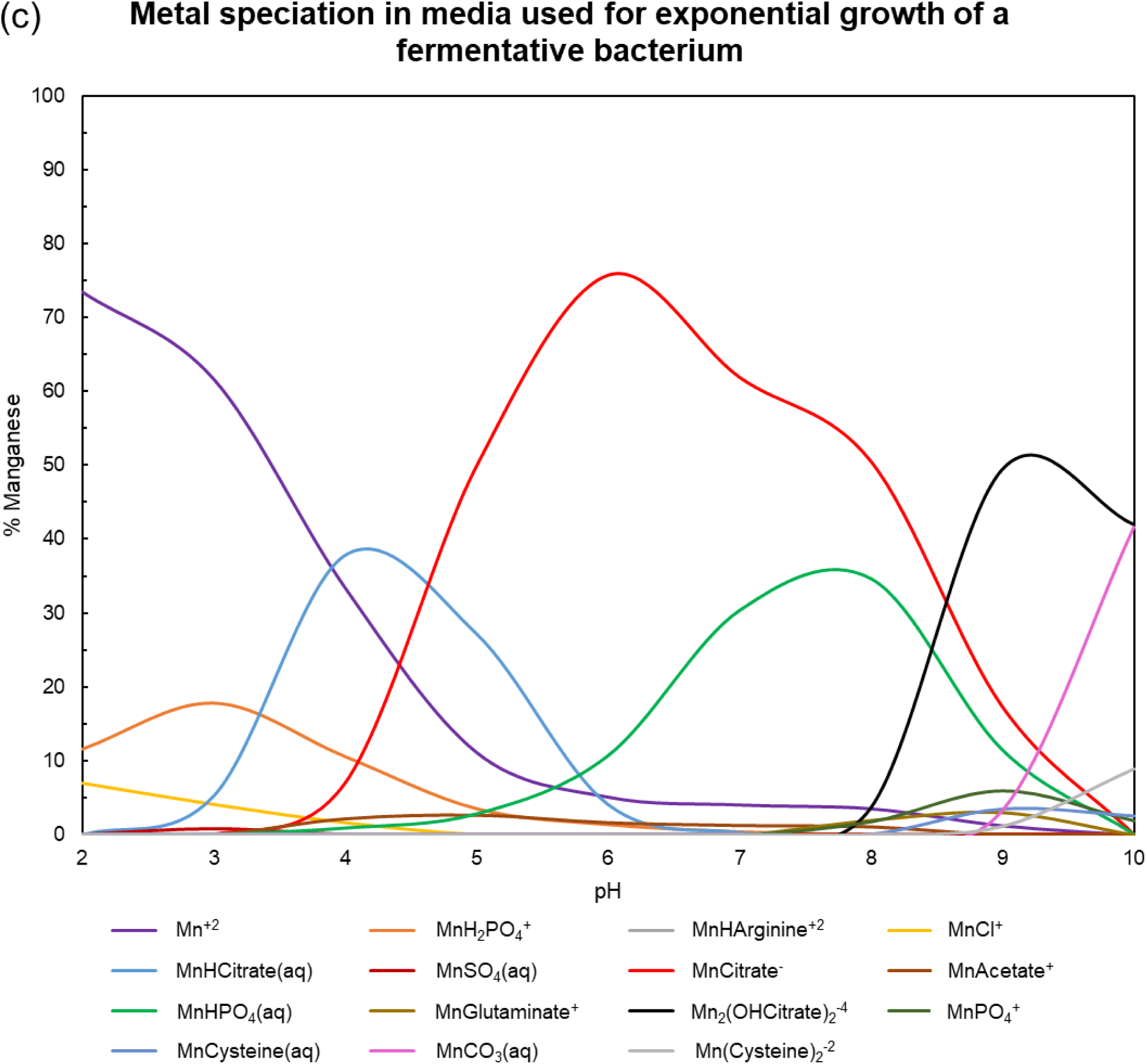
**Manganese speciation in media employed by Kim et al. 2012.**

To illustrate the effect of pH on the ‘stimulatory’ nutrient manganese, we performed speciation calculations from pH of 2 to 10 (Fig. 4c). Greater than 1% of manganese was found to be distributed across 15 species. Expectedly, free manganese and protonated citrate complexes dominated the speciation at low pH while carbonate and hydroxylated citrate complexes dominated at higher pH. The diversity of species suggests that manganese bioavailability is likely to be heavily influenced by pH. We hope such calculations are performed in conjunction of experiments and can help guide and optimize the creation of chemically defined media in the future.

## 6. Metal speciation of defined growth media listed on the DSMZ website

The German Collection of Microorganisms and Cell Cultures (generally referred as DSMZ for their German name-Deutsche Sammlung von Mikroorganismen und Zellkulturen) is one of the most comprehensive resources for microbiology. They acquire and maintain a large collection of cell lines of all taxonomical variety and their website (https://www.dsmz.de/) contains a large list of standardized growth media that are often used to grow microorganisms (Zhang et al. 2018, Zhu et al. 2018, Ghavipanjeh et al. 2018, Haddad et al. 2014). As a service to future scientists that may be interested in the impact of metals in microbial growth, we performed speciation calculations in all chemically defined growth media listed on the DSMZ website. Since the exact chemical compositions of growth media with tryptone, beef extract and yeast extract are not known, speciation calculations could not be performed for these cases.

Out of the 1703 media listed on the DSMZ website, we only found 26 that had exact chemical compositions (chemically defined media). Many of these were variants of a common set of chemicals with minor variations. We obtained metal speciation for six representative media using experimental equilibrium constants reported in the literature or estimated metal-ligand stability constants (Prasad & Shock 2025a, Prasad & Shock 2025b). Speciation for all metals present in these media is shown in Fig. 5a.

**Fig. 5a.**
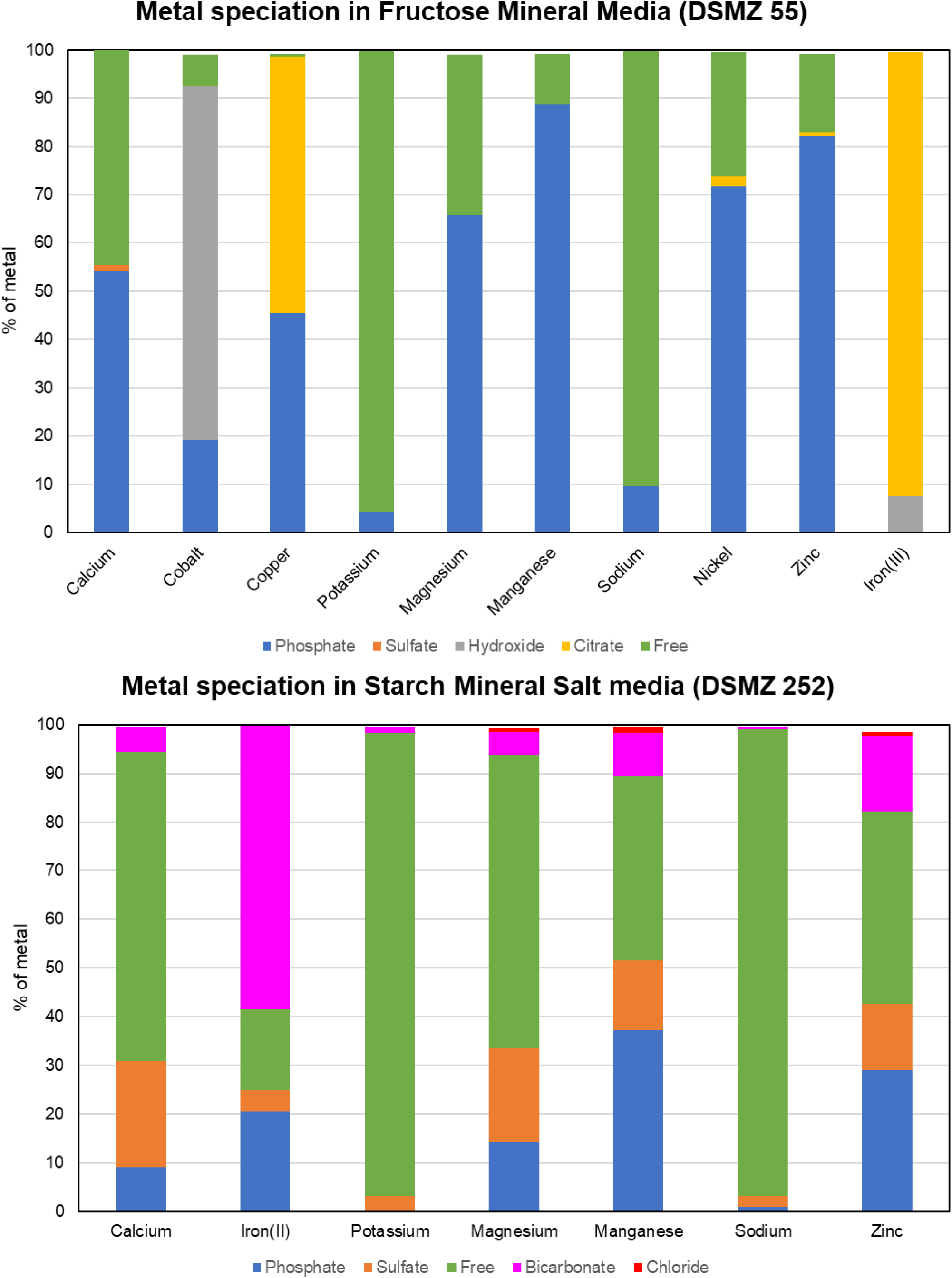
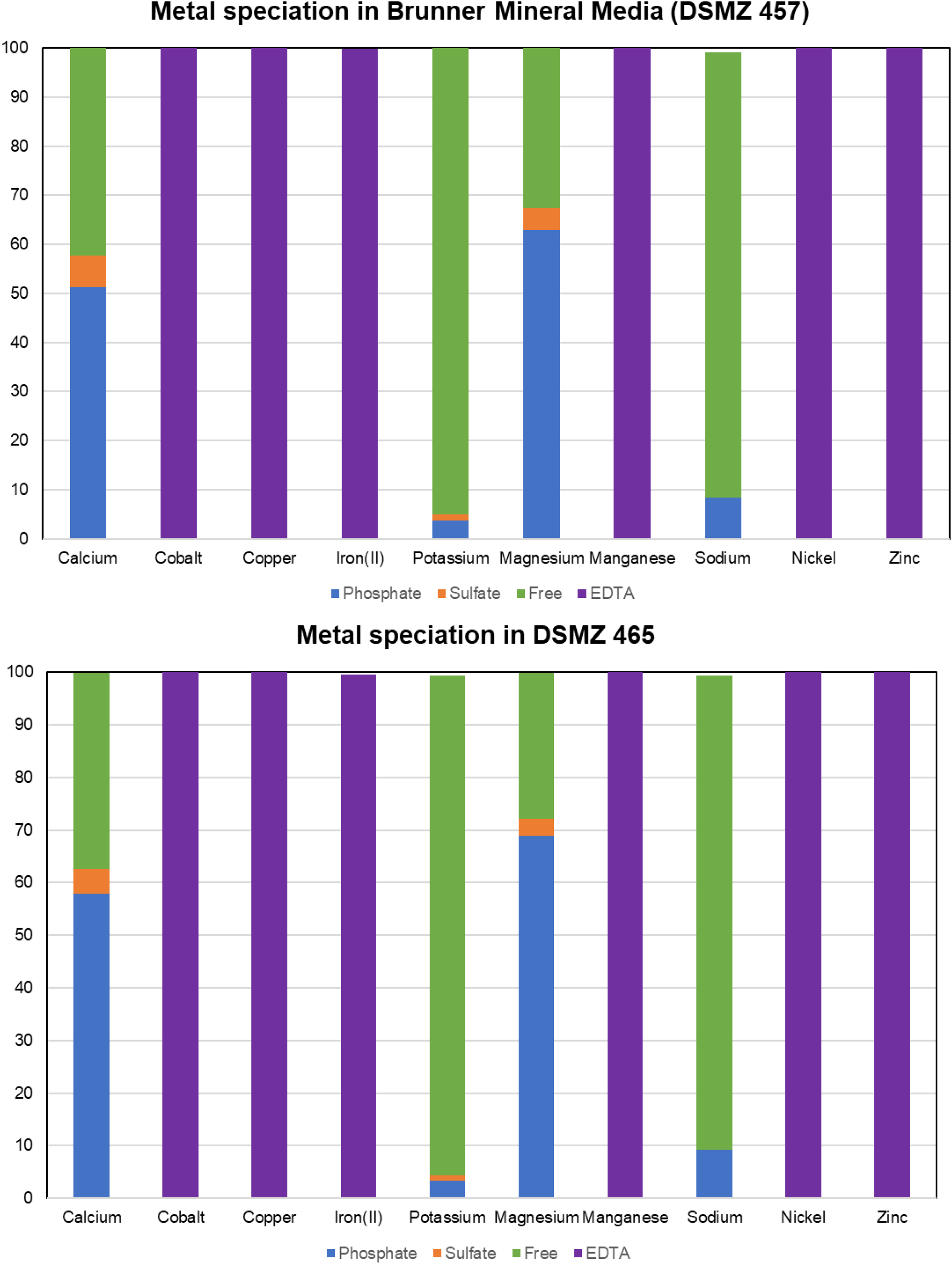
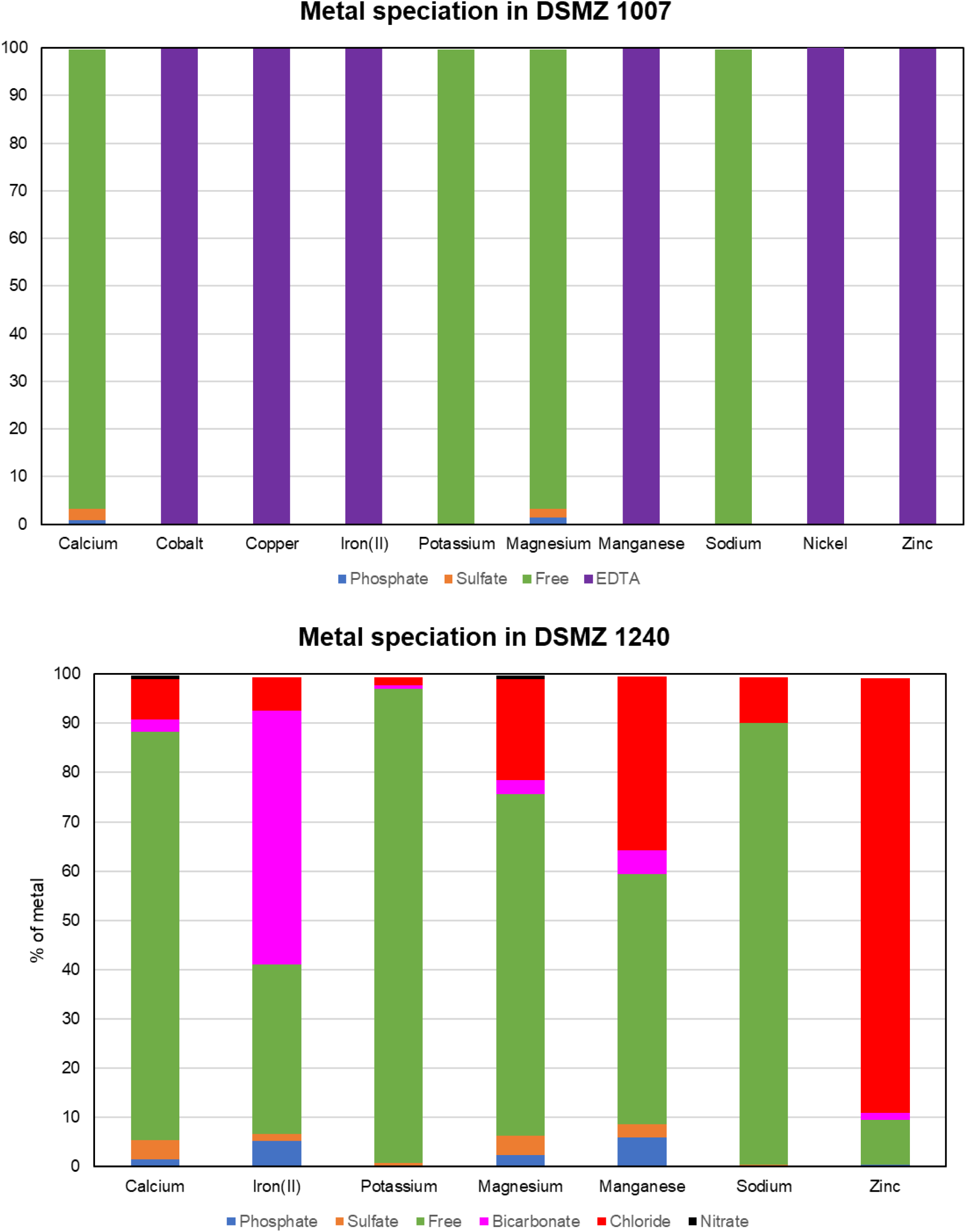
**Metal speciaton of defined growth media at DSMZ website.**

As may be seen from the figure, speciation may be similar or drastically different for metals in the same media. As an illustration, nickel and zinc have similar distribution in the fructose mineral media while cobalt and copper are speciated completely differently. It may be worth noting that all the metals above are divalent transition metals. Other speciation observations of significance are that EDTA dominates the transition metal speciation in DSMZ 457, 465 & 1007 unlike other transition metals and that chloride dominates zinc speciation in DSMZ 1240.

Many of the above chemically defined media like DSMZ 457 and 465, have variants with minor additions to the original composition. For example, DSMZ 457a can be synthesized by adding ∼5mM 2-chlorobenzoate that may be used as a sole carbon and energy source. As 2-chlorobenzoate has a metallophilic carboxylate group, we suspected if this ligand would disrupt the metal speciation in the medium. To theoretically test this, we performed metal speciation in DSMZ 457a using stability constants for metal complexes of 2-chlorobenzoate estimated by Prasad & Shock 2025a. These estimations were made using numerous linear free energy relationships as no stability constant measurements were available for any metal ion. As may be seen from Fig. 5b, the metal speciation in DSMZ 457a is almost identical to DSMZ 457. To investigate the impact of composition on the metal speciation in the medium, we performed these calculations with minor modifications. The metal speciation was largely similar when pH was decreased and increased by 1 unit. Raising the concentration of 2-chlorobenzoate by 10 times changed the speciation of calcium, potassium, magnesium and sodium by a few percent but no change was made to the transition metal speciation. However, reducing EDTA concentration by a factor of 10 significantly changed the speciation of the transition metals iron (II) and manganese. These metals were predominantly distributed as phosphate complexes (mainly HPO_4_^-2^) in this composition with EDTA complexes accounting for less than 6% of the total metal. Speciation for cobalt and zinc was minorly affected as more than 60% of these metals were distributed as their EDTA complexes. On the other hand, copper and nickel distribution was completely unaffected with EDTA chelates predominating the respective speciation. This speciation was largely preserved when the concentration of 2-chlorobenzoate was additionally increased by a factor of 10. Thus, decreasing micromolar EDTA concentration has a bigger impact on metal speciation than increasing millimolar 2-chlorobenzoate concentration. Similar calculations can be performed for many chemically defined growth media using stability constant values estimated in Prasad & Shock 2025a and Prasad & Shock 2025b.

**Fig. 5b.**
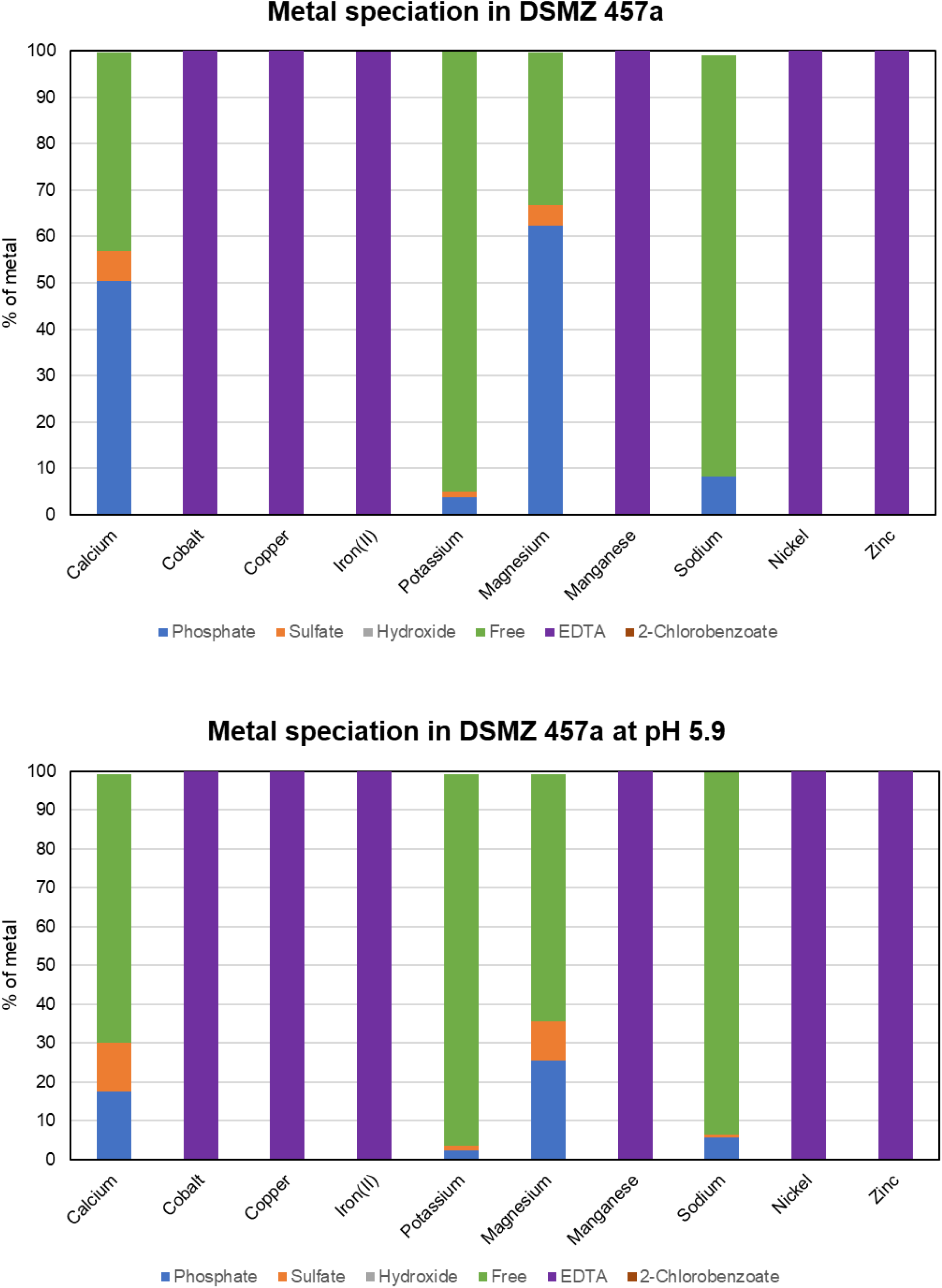
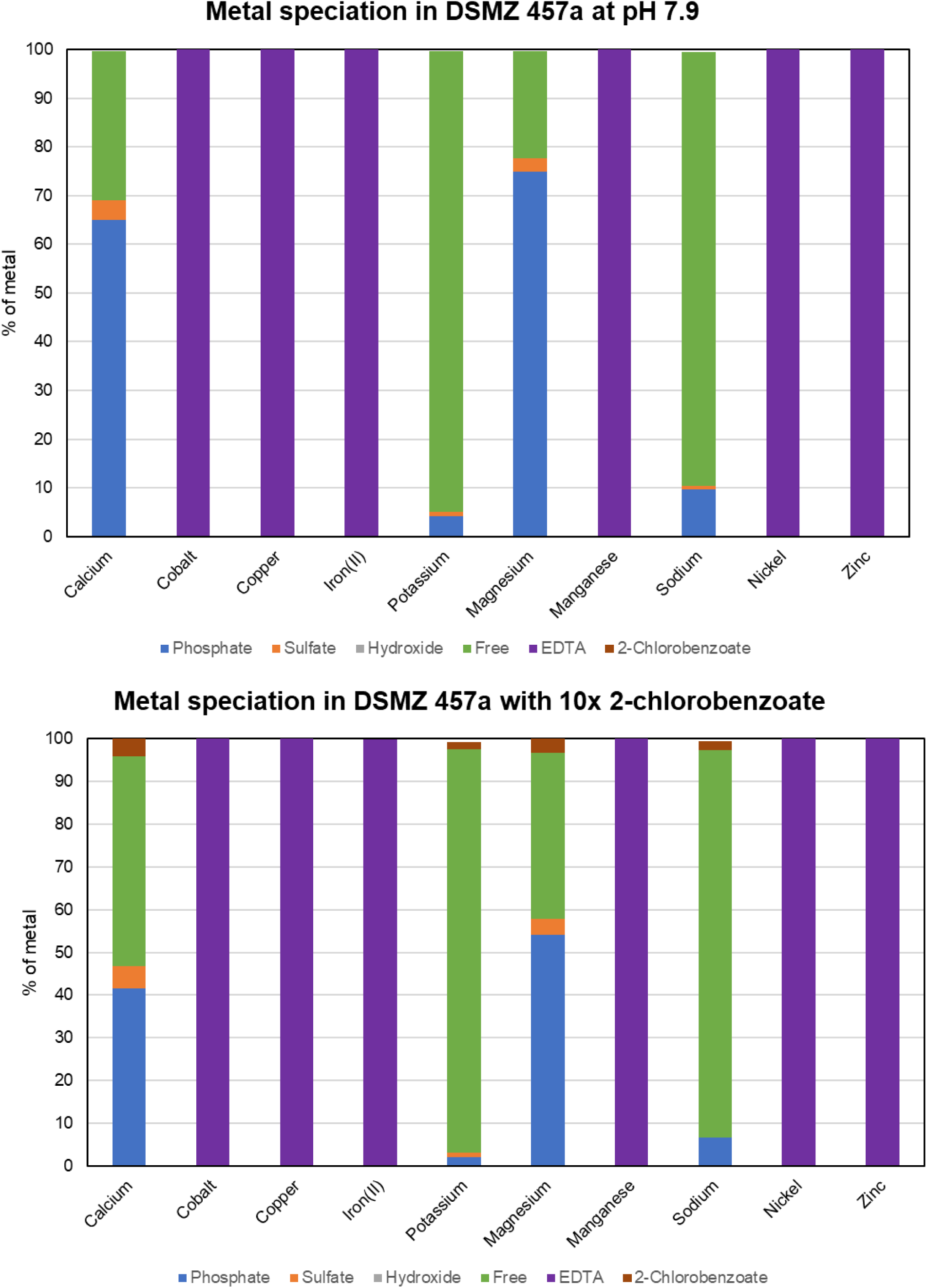
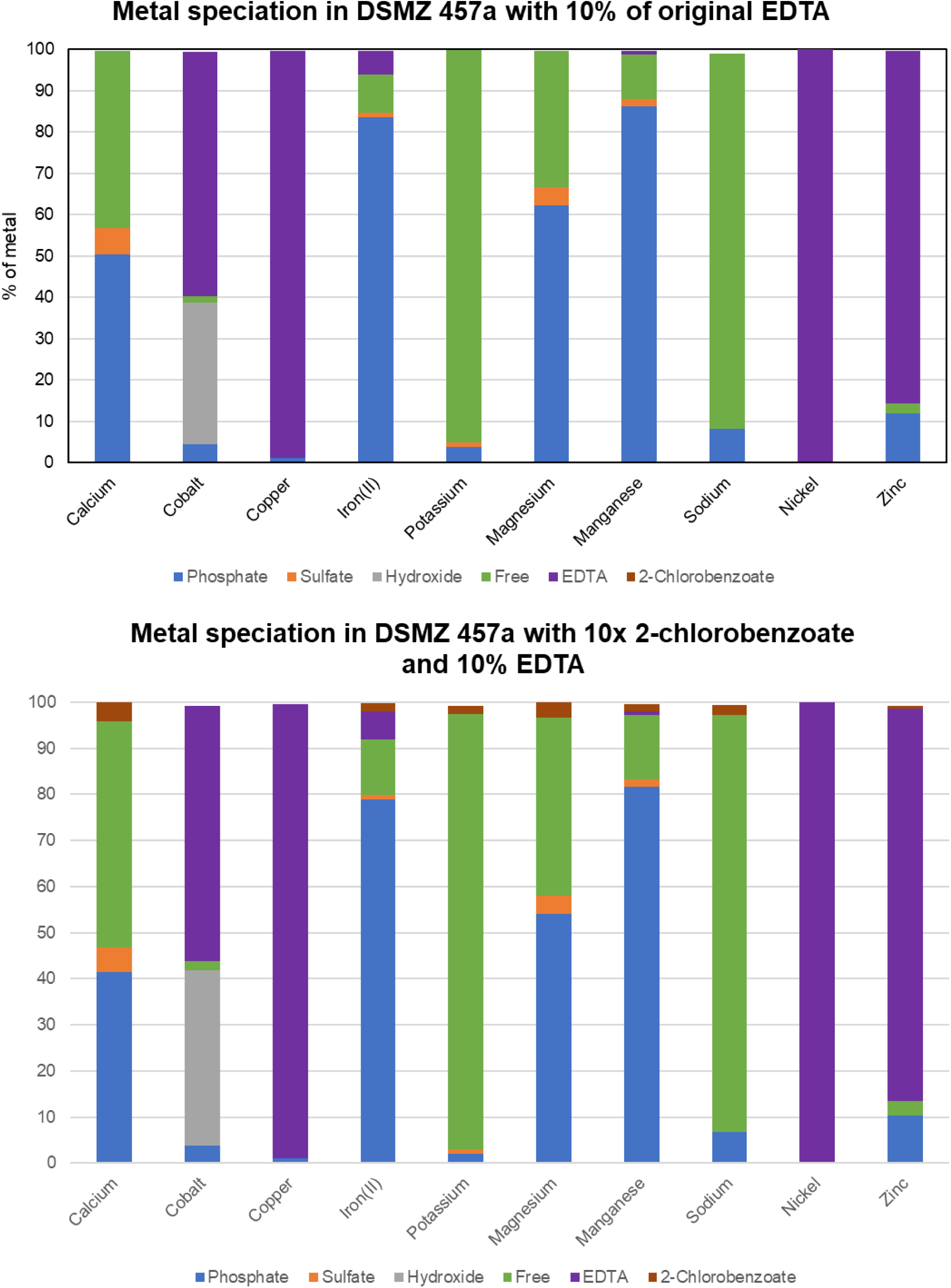
**Metal speciaton in DSMZ 457a with minor variations in composition**

## 7. Conclusion

We have obtained metal speciation in some commonly used microbial growth media employing an updated thermodynamic database. We have evaluated how metal speciation changes with variations in medium composition. Our results suggest that the slightest modifications in pH and chelator concentration can significantly alter metal speciation. Such calculations are only limited by the availability of metal-ligand stability constants. In the case that experimental values for such stability constants were not present in the literature, estimation methods were utilized to fulfill this requirement. We have additionally made correlations of metal speciation with experimental measurements of biological response. These correlations informed us about the metal species taken up by different life forms and the associated uptake mechanisms. We evaluated the applicability of the free ion activity model in two case-studies and critiqued the findings of the original authors.

While a relationship between metal speciation and metal bioavailability was first proposed over 45 years ago, there is still a lot to learn on this topic. A deep understanding of this subject is needed to assess the degree of metal-requirement and the dangers of metal-toxicity to microbes and human cells alike. We hope future microbial growth experiments are performed in conjunction with these speciation calculations that are free of cost and easy to obtain.

## Supporting information

https://data.mendeley.com/datasets/4rv5vfbgcg/1

