## Supplementary material for "Metal speciation and bioavailability in microbial growth media": https://data.mendeley.com/datasets/4rv5vfbgcg/1

### S1 - Comparison of metal speciation calculations with corresponding measurements from a seminal microbial study

In the 1976, Sunda & Guillard published the first study correlating metal speciation and bioavailability (Sunda & Guillard 1976) that eventually gave birth to the free ion activity model (Morel et al. 1978). In the study, the original authors performed copper toxicity experiments for a marine algae in a defined growth medium. They varied the free copper activity by primary altering pH and ligand concentration and measured algal growth rate. Owing to the compositional complexity of the growth medium, the free copper ion activity could not be measured using ion-selective electrodes and were instead obtained using equilibrium speciation calculations. They verified the validity of their calculations by comparing their calculated activity of free copper to copper activity measurements in a simple 'test medium' primarily composed of water, copper and the chelator Tris. We performed a similar comparison in Fig. B1 where the negative logarithm of free copper activity (pCu= -log $a_{Cu+2}$) was calculated using the equilibrium speciation software EQ3/6 (Wolery 2010). As may be seen from the figure, our simulations are in excellent agreement with the corresponding measurements performed using ion-selective electrodes. The agreement is greater for high free copper activity (low pCu value) compared to low free copper activity (high pCu value) that is understandable owing to higher experimental uncertainty at lower free copper concentration. These results encouraged us to simulate other studies using our thermodynamic database and software.

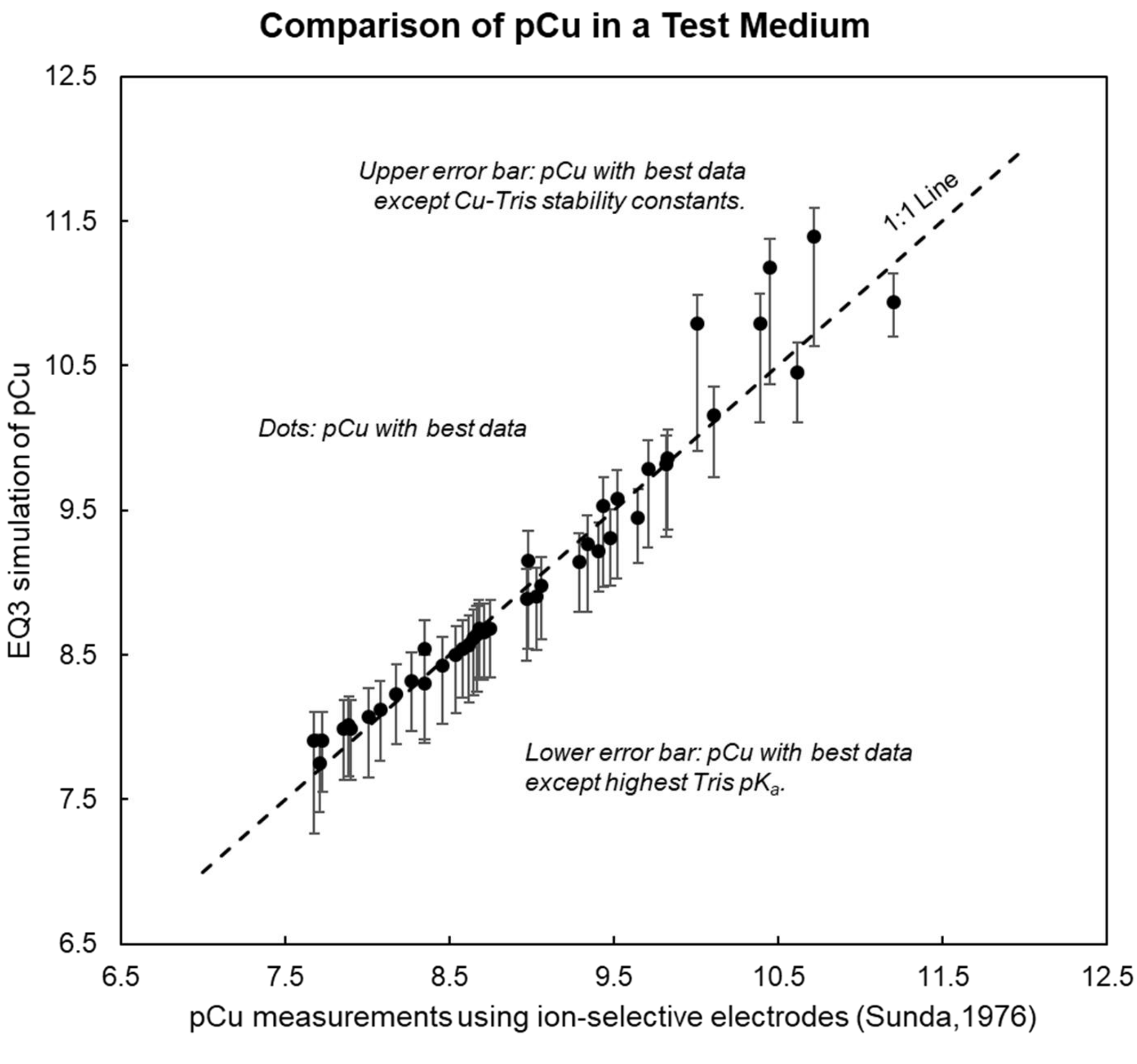


Fig. B1. Comparison of EQ3 calculation of pCu (pCu= -log $a_{Cu+2}$) with experimental measurements made by Sunda & Guillard 1976. Copper-Tris complexes accounted for over 99% of copper's speciation.
